## Supplementary figures and images for "Screening conditions and constructs for attempted genetic transformation of *C. elegans* by *Agrobacterium*"

### Supplemental Table 1

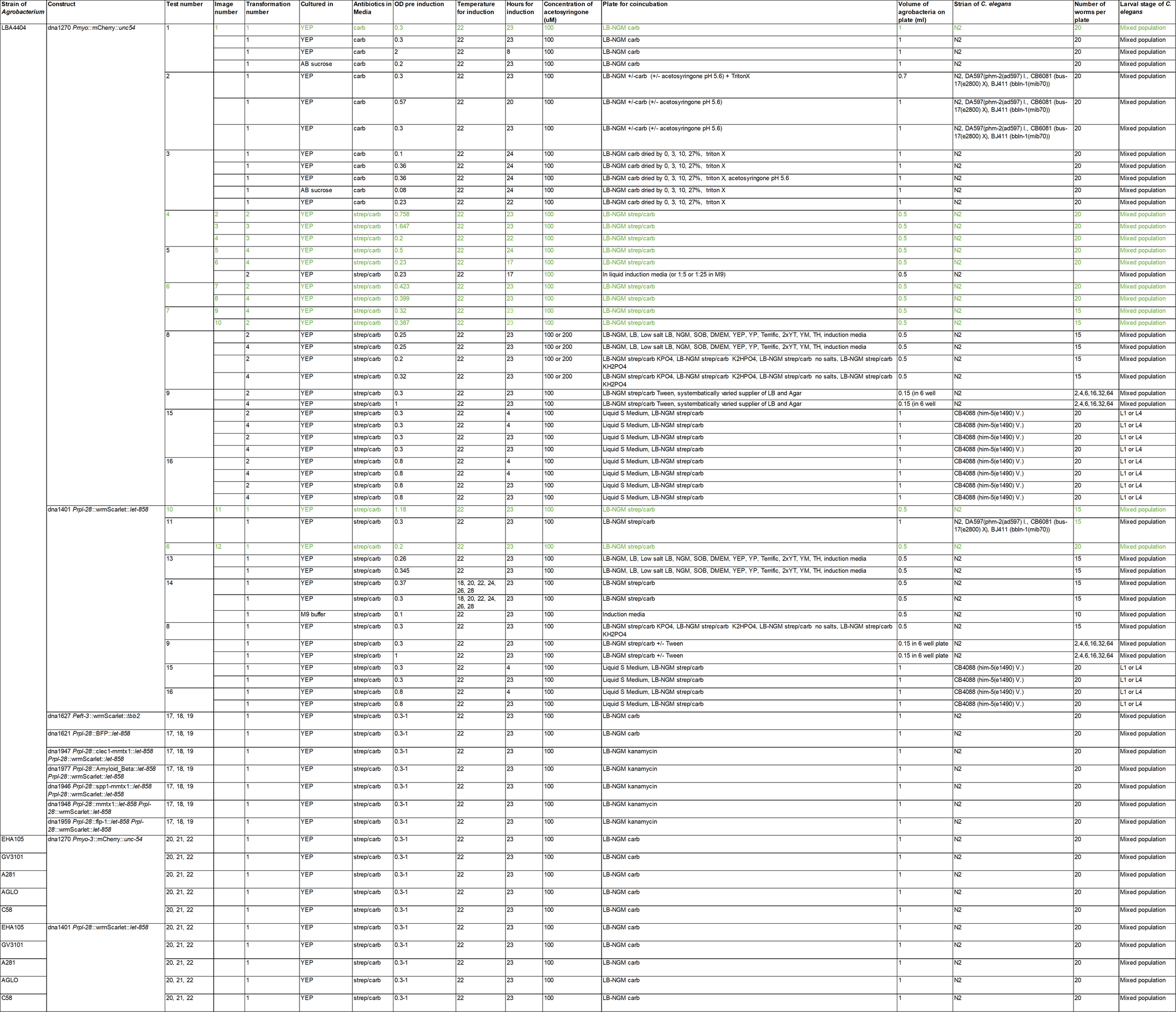

### Supplementary Fig 2

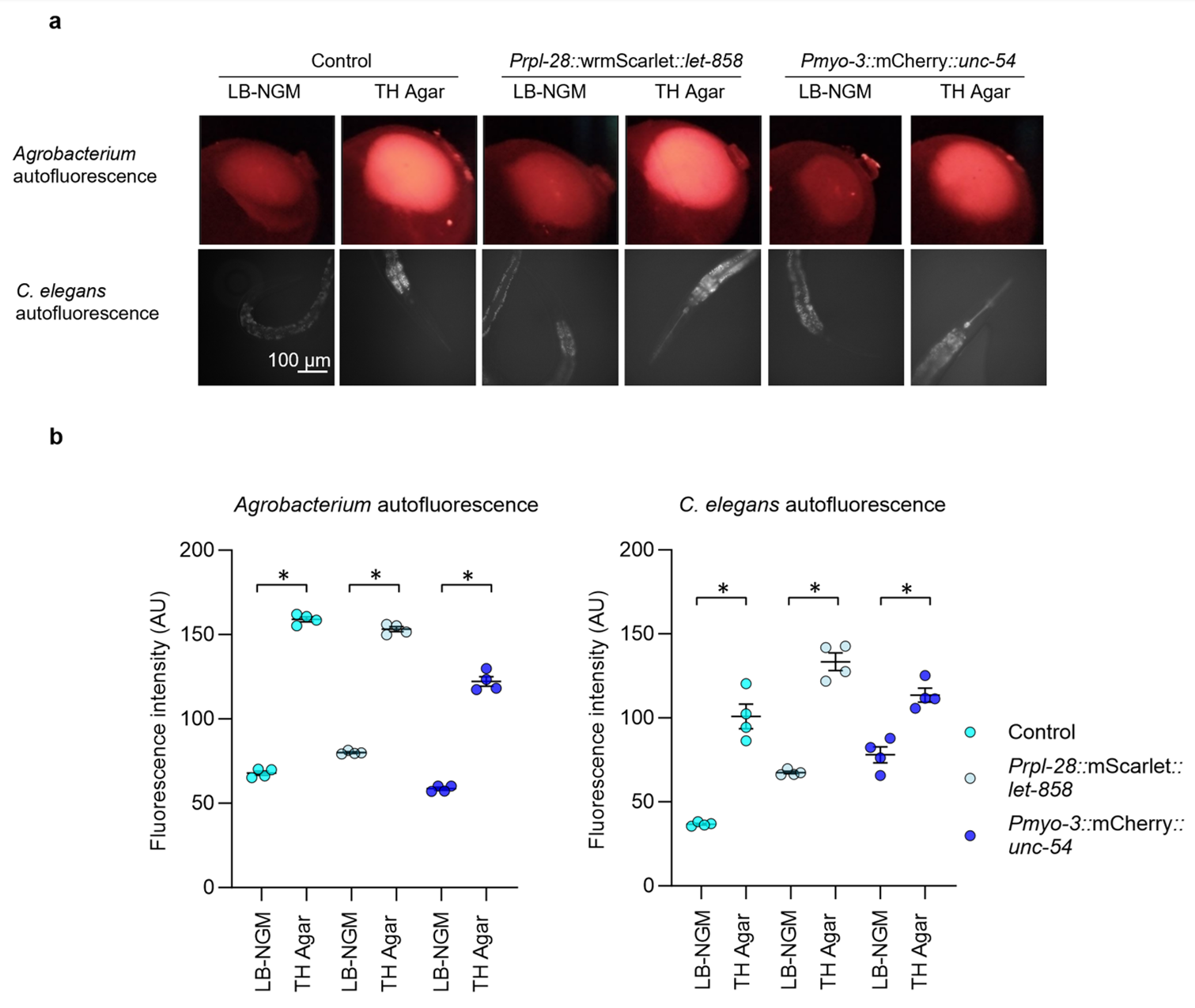

### Supplementary Fig 3

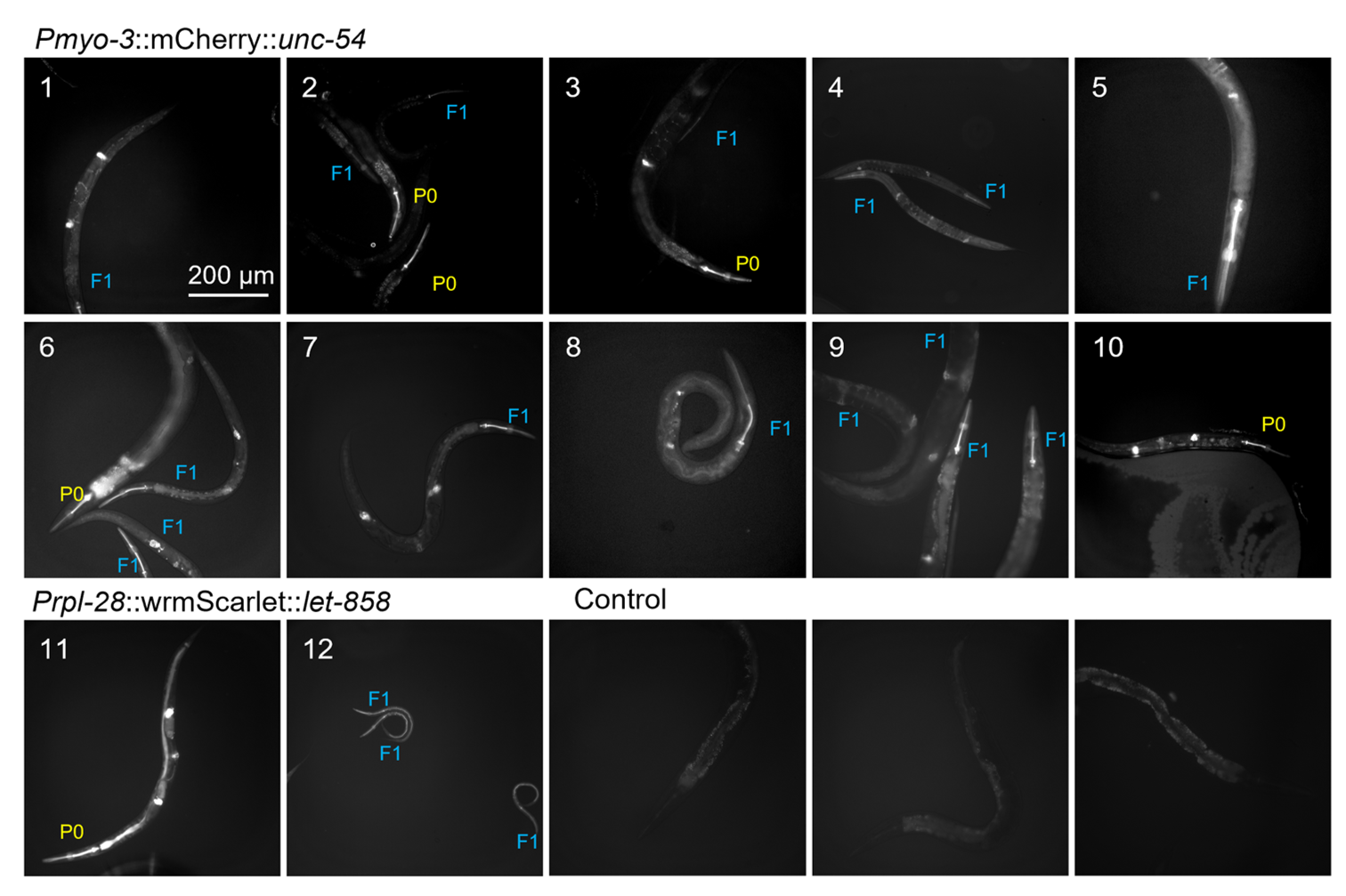

### Supplementary Fig 4

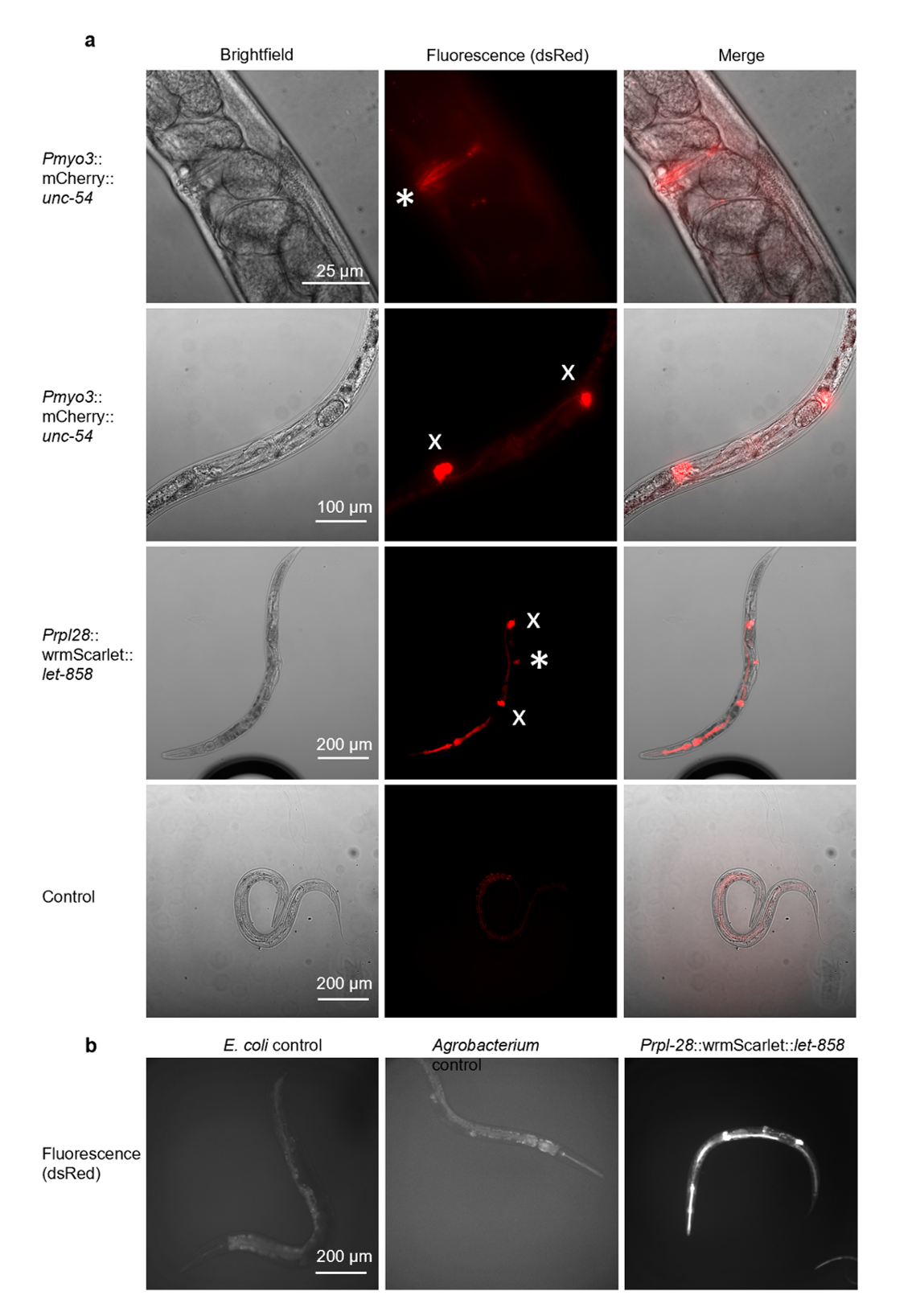

### Supplementary Fig 5

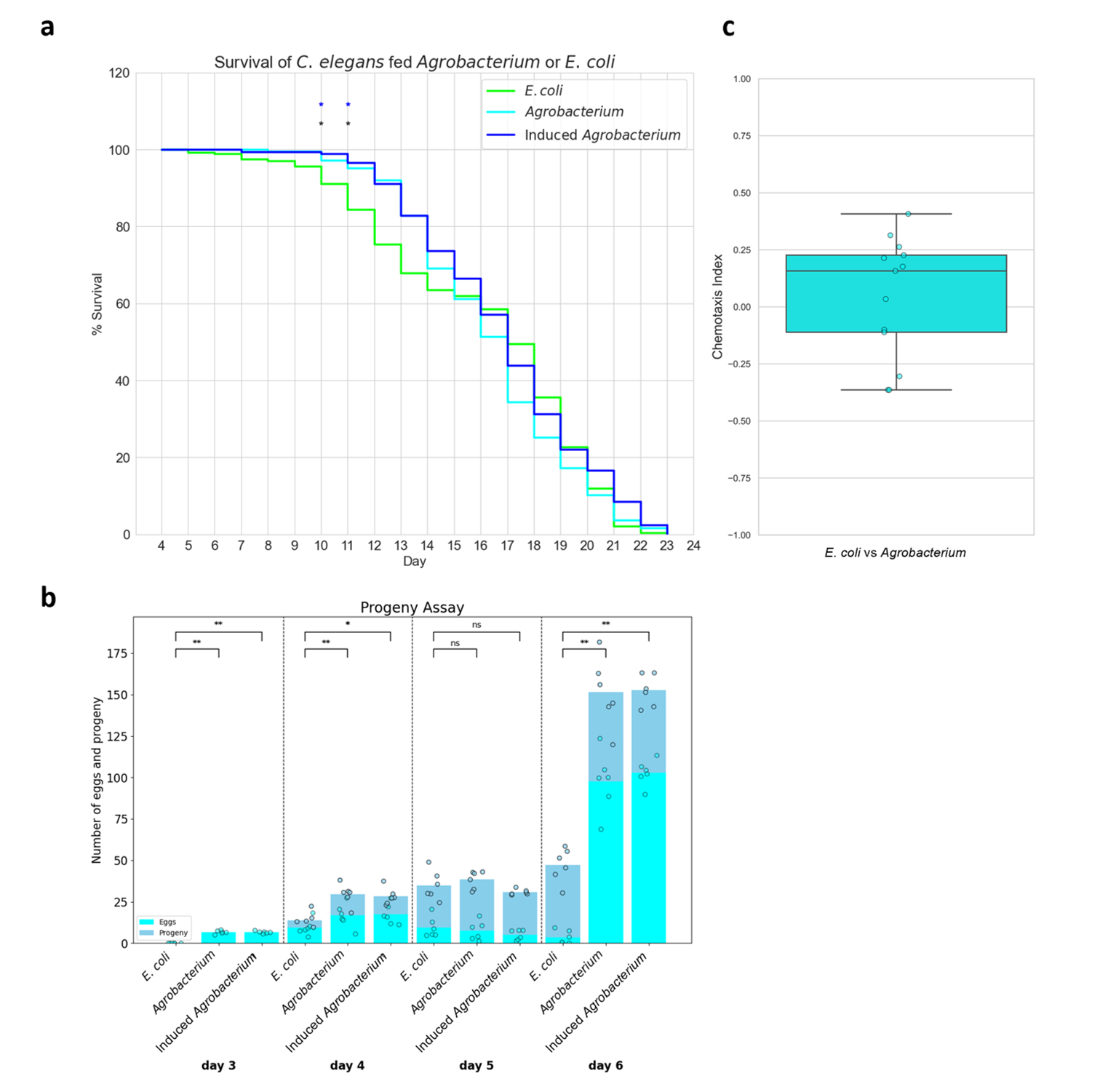
